# VelcroVax: a ‘bolt-on’ vaccine platform technology improves antibody titres against a viral glycoprotein in mice

**DOI:** 10.1101/2022.04.22.489148

**Authors:** Natalie J Kingston, Keith Grehan, Joseph S Snowden, Mark Hassall, Jehad Alzahrani, Guido C Paesen, Lee Sherry, Connor Hayward, Amy Roe, Sam Stephen, Darren Tomlinson, Antra Zeltina, Katie J Doores, Neil A Ranson, Martin Stacey, Mark Page, Nicola J Rose, Thomas A Bowden, David J Rowlands, Nicola J Stonehouse

**Affiliations:** Astbury Centre for Structural Molecular Biology, School of Molecular & Cellular Biology, Faculty of Biological Sciences, University of Leeds, Leeds, LS2 9JT, United Kingdom; Division of Virology, National Institute for Biological Standards and Control (NIBSC), Blanche Lane, South Mimms, Potters Bar, Hertfordshire, EN6 3QG, UK; Division of Structural Biology, Wellcome Centre for Human Genetics, University of Oxford, Oxford, OX3 7BN, UK; Bio Strategy Unit, Boehringer Ingelheim Pharma GmbH, Birkendorfer Strasse 65, 88400 Biberach an der Riss, Germany; Department of Infectious Diseases, School of Immunology and Microbial Sciences, King’s College London, UK

## Abstract

Having varied approaches to the design and manufacture of vaccines is critical in being able to respond to worldwide needs and to newly emerging pathogens. Virus-like particle (VLP) vaccines form the basis of two of the most successful licensed vaccines (against hepatitis B virus (HBV) and human papillomavirus). They are produced by recombinant expression of viral structural proteins, which self-assemble into immunogenic nanoparticles. VLPs can also be modified to present unrelated antigens, and here we describe a universal ‘bolt-on’ vaccine platform (termed VelcroVax) where the capturing VLP and the target antigen (hapten) are produced separately. We utilise a modified HBV core (HBcAg) VLP, with surface expression of a high-affinity binding sequence (Affimer) directed against a SUMO tag and use this to capture SUMO-tagged gp1 glycoprotein from the arenavirus, Junín virus (JUNV). Using this model system, we have solved high-resolution structures of VelcroVax VLPs, and shown that the VelcroVax-JUNV gp1 complex induces superior humoral immune responses compared to the non-complexed viral protein. We propose that this system could be modified to present a range of haptens and therefore form the foundation of future rapid-response vaccination strategies.

## Introduction

The need for safe and effective vaccines to be developed rapidly and distributed globally has been highlighted over the last two years. Vaccines have been developed for more than 20 different pathogens, and more than 15 additional organisms are recognised by the World Health Organization (WHO) as priority pathogens with epidemic or pandemic potential. Although the WHO endeavours to accelerate the development of vaccines for these priority pathogens for use in low- and middle-income countries (LMICs), there are significant challenges to their development and deployment^1^. These include safety, efficacy and the need to maintain a cold chain when delivering vaccines to remote areas. Importantly, the availability of vaccines in endemic regions is essential to control the spread of pathogens and facilitate the prevention of future global pandemics.

The list of pathogens with epidemic or pandemic potential varies among global authorities. The National Institute of Allergy and Infectious Disease (NIAID) priority list includes some new world arenaviruses, including Junín virus (JUNV), which causes a potentially lethal haemorrhagic disease known as Argentine haemorrhagic fever (AHF)^2,3^. JUNV is transmitted via rodents *(Calomys musculinus)* and contracted via contact with infected excretions or aerosols. Outbreaks of AHF in the 1960s and 1970s resulted in thousands of deaths and had case fatality rates between 15-30%^4–6^. Total cases decreased in the following decades and have fallen substantially since the introduction of a live attenuated vaccine in affected regions of Argentina^6–8^. Despite the success of this vaccine, as with all attenuated virus vaccines, there remain safety concerns regarding the potential for reversion to a pathogenic form.

The advancement of technologies used for vaccine production and purification have contributed to the generation of safer vaccines. For example, virus-like particle (VLP) vaccines for hepatitis B virus (HBV) and human papillomavirus (HPV) have shown exemplary safety and efficacy^9–10^. Most recently, recombinant non-replicating viral vectors and RNA vaccines have been produced rapidly and also show impressive safety and efficacy profiles^11–13^. Critically, in contrast to attenuated vaccines, inactivated, subunit, polysaccharide, RNA or toxoid vaccines are non-replicative, so do not pose the risk of reversion to a pathogenic form. This makes recombinant technologies the most attractive approach for the development of safer next-generation vaccines.

The efficacy of subunit vaccines can be enhanced when the subunit exists as a nanoparticle. Nanoparticles may be naturally occurring (VLPs), artificially formed^14–16^ or modified biological chimeras^17–19^. Indeed, chimeric VLP technology has allowed the deployment of the first licensed anti-malaria vaccine, Mosquirix^18,20^. The success of this vaccine suggests that a chimeric VLP approach is both tractable and suitable for improving responses against challenging immunogens, however, it took over 30 years for Mosquirix to be licensed^18,21^. Alternative approaches for modifying VLPs have been investigated to increase the diversity of vaccine platforms. The approach we have pursued relies upon the surface display of a capturing molecule (e.g. antibody, nanobody) on a nanoparticle carrier, which is able to bind and display an antigen of interest (hapten) tagged with an appropriate sequence. Poorly immunogenic haptens displayed on nanoparticles are more effectively recognised by dendritic cells (DCs). In addition, nanoparticle size (30-100 nm) can influence T helper bias and T helper epitopes from protein-based nanoparticles can contribute to anti-hapten immunity, thus humoral responses generated are likely to be higher affinity and more diverse^22–26^.

Here, we describe a vaccine system in which a carrier nanoparticle and hapten are produced separately. We have developed a modified HBV core (HBcAg) VLP, termed VelcroVax, with surface expression of a SUMO-Affimer. Affimers are produced by phage display approaches, are small (~13 kDa) and can be expressed in a range of systems^27^. We used these VLPs to capture the SUMO-tagged JUNV glycoprotein, gp1. Using this model system, we characterise VelcroVax structurally and functionally, using comparative immunisation trials to determine whether JUNV gp1 coupling to VLPs alters the immune response generated. We propose that this system may be modified for a range of haptens and could form the foundation of future rapid-response vaccination strategies.

## Methods

### Generation of HBcAg VLPs in yeast

Genes encoding either HBcAg or VelcroVax were introduced downstream of the AOX1 promoter within the pPinkHC expression vector (ThermoFisher Scientific). The VelcroVax sequence consists of a fused HBcAg dimer with the SUMO-Affimer sequence introduced within the first major immunodominant region (MIR) of this dimer. A Gly-Ser linking sequence was used to provide flexibility to this domain, and for consistency this Gly-Ser linker was present in all HBcAg subunits used here. Similar to previously described protocols^28^, plasmids were linearised with *AflII* and electroporated into PichiaPink strain 1 (Invitrogen), then transformed yeast were plated on adenine dropout (PAD) media and incubated at 28°C for 3-5 days. To screen for expression, colonies were selected at random and inoculated into 5 mL YPD media (10 g/L yeast extract, 5 g/L peptone, 20 g/L Dextrose) before incubation at 28°C, 250 rpm for 48 hours. Cells were pelleted at 1,500 rcf and resuspended in 1 mL YPM (10 g/L yeast extract, 5 g/L peptone, 2% v/v methanol). Cultures were incubated at 28°C, 250 rpm for 72 hours, and supplemented with 1 or 2% v/v methanol every 24 hours (VelcroVax and HBcAg expressions, respectively). Cells were collected at 48 hours and assessed for scFv production by western blot.

For large-scale production, a glycerol stock of VelcroVax- or HBcAg-expressing *P. pastoris* was used to inoculate 5 mL YPD and incubated at 28°C for 48 hours at 250 rpm before inoculation into 200 mL of YPD and incubation for a further 48 hours at 28°C, 250 rpm. Cells were pelleted at 1,500 rcf and resuspended in 200 mL YPM (1 or 2% v/v methanol, as above) before incubation at 28°C, 250 rpm for 72 hours. Media were supplemented with methanol every 24 hours. Cells were pelleted at 4,000 rcf and resuspended in 30 mL EDTA-free breaking buffer (50 mM Na_3_PO_4_, 5% v/v glycerol, pH 7.4) with cOmplete EDTA-free protease inhibitor cocktail (Roche).

### VLP purification and quantitation

To isolate VLPs from *P. pastoris*, cells were disrupted at 40 kpsi and supplemented with 1 mM MgCl and 250 units denarase (c-LEcta) before incubation at room temperature for 2 hours with agitation. Samples were clarified at 4,000 rcf and clarified supernatant was precipitated overnight at 4°C with 20% v/v saturated ammonium sulphate solution (structural studies) or 8% w/v PEG-8000 (immunogenicity and antigenicity studies). Precipitated material was pelleted at 4,000 rcf for 30 minutes and re-suspended in 30 mL PBS. Insoluble material was removed by centrifugation at 10,000 rcf. The soluble material was pelleted through a 30% sucrose cushion at 150,000 rcf for 3.5 hours. Pellets were resuspended in 1 mL PBS and separated on a 15-45% sucrose gradient at 50,000 rcf for 12 hours. 1 mL fractions were collected manually (top down) and assessed for the presence of HBcAg-reactive proteins by western blot with mAb 10E11 using standard protocols. The protein content of fractions was assessed directly by BCA assay (Pierce, ThermoFisher Scientific), or the VLPs were concentrated, and buffer exchanged using 100k mwco PES concentrator columns (Pierce, Thermo Scientific) before quantification by BCA assay. To purify VLPs for structural analysis, this protocol was slightly modified, as described in Snowden et al. (2021)^29^.

### Electron microscopy

To prepare samples for negative stain EM, carbon-coated 300-mesh copper grids (Agar Scientific, UK) were glow-discharged under air (10 mA, 30 s) before applying 3 μL sample for 30 s. Excess liquid was wicked away, then grids were washed two to four times with 10 μL distilled H2O. Staining was then performed with 1 – 2% uranyl acetate solution (UA). UA was applied (10 μL) and immediately wicked away, then an additional 10 μL UA was applied and allowed to incubate for 30 s prior to blotting and leaving to air dry. Imaging was performed using either (i) an FEI Tecnai G2-spirit with LaB_6_ electron source, operating at 120 kV and equipped with a Gatan Ultra Scan 4000 CCD camera, with a calibrated object sampling of 0.48 nm/pixel, or (ii) an FEI Tecnai F20 with field emission gun, operating at 200 kV and equipped with an FEI CETA camera, with a calibrated object sampling of 0.418 nm/pixel.

For cryoEM, samples were vitrified as described in Snowden et al (2021)^29^. Briefly, ultrathin continuous carbon-coated lacey carbon 400-mesh copper grids (Agar Scientific, UK) were glow discharged in air (10 mA, 30 s), then 3 μL sample were applied to the grid surface for 30 s in a humidity-controlled chamber (8°C, 80% relative humidity). Excess liquid was removed by blotting (1.0 – 4.0 s) before plunge freezing in liquid nitrogen-cooled liquid ethane using a LEICA EM GP plunge freezing device (Leica Microsystems, Germany). Imaging was performed using an FEI Titan Krios transmission EM (ABSL, University of Leeds) operating at 300 kV, with a calibrated object sampling of 1.065 Å/pixel. Full data collection parameters are provided in Table S1.

### Image processing

Image processing was performed using the Relion 3.0 and Relion 3.1 pipelines^30^,^31^. MotionCor2^32^ was used to correct any motion-induced blurring in raw micrographs, then CTF parameters were estimated using Gctf^33^. A small subset of VLPs (both *T* = 4 and *T* = 3*) was manually selected and used to generate 2D class averages, used as templates for automated picking of the entire dataset. Initially, ~250,000 particles (including contaminants and erroneously selected areas of carbon) were extracted and 2× down-sampled for 2D classification, with CTFs ignored until the first peak. All classes resembling VLPs (~130,000 particles) were taken forward for additional 2D classification without CTFs ignored until the first peak, at which point two independent particle stacks were created and re-extracted without down-sampling: one for *T* = 4 VLPs and one for *T* = 3* VLPs (each containing ~50,000 particles). 3D refinement was performed separately for each particle stack, based on initial models generated *de novo* in Relion, with symmetry imposed (I1 for *T* = 4, C5 for *T* = 3*). Where appropriate, map resolution and quality were improved by iterative cycles of CTF refinement, Bayesian polishing and 3D refinement with a solvent mask applied and flattened Fourier shell correlation (FSC) calculations. Maps were sharpened using a solvent-excluding mask and a nominal resolution determined using the ‘gold standard’ FSC criterion (FSC = 0.143) (Figure S2, Table S2), then local resolution was calculated and a local resolution-filtered map generated in Relion.

For *T* = 4 VLPs, focussed classification was performed in an attempt to resolved Affimer density, using a protocol described previously^29,34–36^. Briefly, SPIDER^37^ was used to generate a cylindrical mask which was manually placed above a four-helix bundle using UCSF Chimera^38^. A soft-edge was added to the mask in Relion. *T* = 4 VLP particles and their associated orientational information from a symmetrised 3D refinement were used to generate a symmetry-expanded particle stack using the relion_symmetry_expand tool. This data was then subjected to masked 3D classification without alignments, with a regularisation parameter of 40.

### Model building and refinement

Atomic models were built into the density maps for both *T* = 4 and *T* = 3* VLPs. Firstly, a homology model was generated using SWISS-MODEL^39^. Copies of this model were fitted into density for each quasi-equivalent position within the *T* = 4 and *T* = 3* VLP asymmetric units using UCSF Chimera^38^, and unresolved segments of the peptide backbone were removed. Models were then inspected and manually refined in Coot^40^ before automated refinement in Phenix^41^ to improve model-to-map fit and atomic geometry. This process was repeated iteratively, with at least one iteration performed with a symmetrised atomic model to avoid erroneous placement of atomic coordinates in density from adjacent asymmetric units. Model validation (Table S2) was performed using MolProbity^42^.

### Structure analysis and visualisation

Visualisation of structural data was performed in UCSF Chimera^38^, UCSF ChimeraX^43^ and PyMOL (The PyMOL Molecular Graphics System, Version 2.1, Schrödinger, LLC). RMSD calculations were performed using the ‘MatchMaker’ tool in UCSF Chimera, with default settings.

### Generation of recombinant JUNV gp1

The sequence encoding amino acids 87-231 of JUNV gp1 (GenBank ACO52428) was PCR-amplified and cloned into a pHLsec vector^44^ containing a C-terminal SUMO tag (GenBank AVL26008.1) and hexahistidine tag. The JUNV gp1-SUMO construct was transfected into human embryonic kidney (HEK) 293T cells, grown in roller bottles for transient expression^45^. Four days post-transfection, cell supernatant was supplemented with NaCl (700 mM), Tris pH 8.0 (20 mM) and imidazole (15 mM). JUNV gp1 was purified by immobilized metal affinity chromatography, using a 5-mL HisTrap Excel column (Cytiva), followed by size-exclusion chromatography (SEC) with a Superdex 200 increase 10/300 GL column (Cytiva) equilibrated with 15 mM Tris (pH 8.0), 200 mM NaCl, and 0.5 mM EDTA. The JUNV gp1 containing peak was further purified over a 1-mL HiTRAP Q (HP) column (Cytiva) using a 30 mM Tris pH 8.0 running buffer and a linear, 0-500 mM NaCl gradient. The JUNV gp1 was re-purified by SEC (as above). Following concentration, protein samples were snap-frozen and stored at −80 °C.

### ELISA to detect antigen capture

The capture of SUMO-tagged JUNV gp1 by VelcroVax was assessed by ELISA. Plates were coated with 50 μL 2 μg/mL of wt HBcAg VLP, VelcroVax or PBS and incubated overnight at 4°C. Plates were blocked with 2% skim-milk powder in PBS 0.1% Tween-20 and JUNV gp1 was added to wells at 1000, 500 and 250 ng/mL, PBS was used as a negative control. Plates were incubated at 37°C for 1 hour before being washed. The presence of JUNV gp1 was determined using a 1:2000 dilution of mouse anti-JUNV gp1 (obtained through BEI Resources, NIAID, NIH: Monoclonal Anti-Junin Virus, Clone OD01-AA09 (immunoglobulin G, Mouse), NR-2567). After incubation, plates were washed and 50 μL of anti-mouse HRP was added to wells (Sigma). Plates were incubated for a further hour at 37°C, before a final wash step and the addition of 100 μL/well Sigmafast OPD (Sigma). After 15 minutes, 50 μL of 3M HCl was added to wells to stop the reaction and the OD was determined at 492 nm. Data was graphed as mean OD 492 nm with SEM (GraphPad Prism).

### Immunisation

Groups of 7 female BALB/c mice were purchased from Charles River UK at 5 weeks of age. Mice were housed for 2 weeks before the initiation of experimental procedures, at which point a sample of pre-immune sera was collected (approximately 50 μL total blood volume) via the tail vein. Mice were then immunised three times at two-week intervals subcutaneously in the rear upper flank with a total volume of 100 μL per dose. Vaccines were composed of 1 μg of VLP (HBcAg or VelcroVax) and 1 μg of JUNV gp1 in the presence of 2.5 nmol CpG ODN1668 (Invivogen). Samples were assembled 24 hours pre-immunisation to facilitate SUMO-linked conjugation of JUNV gp1 to VLP and stored at 4°C until used. All vaccine components were tested for endotoxin content and immunisations contained less than 2.5 EU/dose (Pierce LAL Chromogenic Endotoxin Quantitation kit, Thermo Scientific). Serum samples were collected on days 13 and 27 (as above) (Fig S5). On day 41 final blood samples were taken via cardiac puncture while mice were euthanised under sodium pentobarbitone. All animals were housed under specific pathogen-free conditions and monitored for wellbeing. All animal procedures were performed in strict accordance with UK Home Office guidelines, under licence PP2876504 granted by the Secretary of State for the Home Office which approved the work described, in accordance with local ethical guidelines and internal committee approval for animal welfare at NIBSC. This study conforms to all relevant ethical regulations for animal work in the UK.

### Antibody titration and isotyping

Antibody titres were assessed by indirect ELISA. To this end, 96-well EIA plates were coated with 50 μL 2 μg/mL target protein. Serum samples were assessed against HBcAg, VelcroVax and JUNV gp1. Plates were blocked with 2% skim-milk powder in PBS 0.1% Tween-20 before the addition of duplicate dilutions of antisera at 1:250-4000 or a PBS-only negative control and incubated at 37°C for 1 hour. Plates were washed and 50 μL of rabbit anti-mouse HRP was added at 1:2000 dilution (Sigma). Plates were incubated at 37°C for 1 hour, washed and incubated with 100 μL Sigmafast OPD (Sigma) for 15 minutes. Reactions were stopped with the addition of 50 μL 3M HCl and optical density read at 492 nm. Data was graphed as a box and whisker plot and the mean OD from PBS-negative wells depicted as a dotted line along the graph for reference (GraphPad Prism).

To determine the isotypes of antibodies generated by immunisation, plates were coated and blocked, as above. Sera were diluted 1:125 and 50 μL of sera or a PBS-only negative control were added to duplicate wells, before incubation at 37°C for 1 hour. Plates were washed and 50 μL of isotype-specific goat anti-mouse antibody was added at 1:1000 dilution (Sigma). Plates were incubated at 37°C for 1 hour before being washed and adding 50 μL of anti-goat HRP (Sigma). After a final 1-hour incubation, plates were washed and developed with OPD, as above. The OD 492 nm of negative control wells (no sera) was deducted from the isotype-specific signal and mean OD graphed on a bar chart (GraphPad Prism).

### Generating pseudovirus

Previously described protocols were used to generate JUNV pseudovirus with minimal modification^46^. Briefly, HEK293T/17 cells were seeded at 30% density and incubated overnight to allow growth to 50-60% confluence at time of transfection. The following day DNA transfections were carried out by combining 1 μg p8.91 plasmid, with 1.5 μg Pcsflw^47^ and 1.5 μg of pCAGGS-JUNV gp in 100 μL of Opti-MEM (Gibco) in a standard microcentrifuge tube, separately 12 μL of 1 mg/mL 25,000 mw linear PEI was diluted in 100 μL of Opti-MEM (Gibco). Tubes were incubated at RT for 5 minutes before PEI mix was added to DNA. The combined mixture was incubated at RT for 15 minutes before being added dropwise to culture media. Plates were incubated for 72 hours at 37°C, 5% CO_2_, at which point media was harvested and filtered through a 0.45 μm PES filter.

### Pseudovirus titration

Harvested cell supernatant containing JUNV pseudovirus was titrated as previously described with minimal modification using RD cells^46^. Briefly, in a 96-well white plate (Greiner Bio-One) 50 μL of pseudovirus-containing supernatant was added per well following a 2-fold serial dilution. Dilutions were added to wells containing 1×10^4^ RD cells/well and incubated for 72 hours at 37°C, 5% CO_2_. The relative luminescence units (RLU) were measured using the Bright-Glo (Promega) luciferase system.

### Neutralisation assay

Triplicate wells of diluted serum and 1×10^5^ RLU of JUNV pseudovirus were added to wells of a 96-well white opaque plate in a final volume of 100 μL. Plates were incubated for 1 h at 37°C, 5% CO_2_ in a humidified incubator, and 1×10^4^ RD cells were added to each well. Plates were incubated for 72 h before RLU was recorded, as above. For 1:100 diluted serum raw data is graphed, for 1:10 diluted samples percentage neutralisation was determined relative to positive (no antibody) and negative (no pseudovirus) wells.

## Results

### Generation of VelcroVax

HBcAg monomers assemble into paired dimers, which further assemble to form *T* = 3 (90 dimers) and *T* = 4 (120 dimers) symmetric particles. Within each dimer the C-terminal end of one monomer is in proximity to the N-terminal end of the other partner (Fig 1A). The genetic fusion of these monomers using a sequence encoding a Gly-Ser linker ensures that the genetically fused pairs will dimerise within the assembled VLP, termed tandem HBcAg (tHBcAg). We inserted a sequence encoding a SUMO-Affimer into the major immunodominant region (MIR) of the first HBcAg monomer within the tandem construct (Fig 1B). This organisation ensures that within each HBcAg dimer, one MIR will contain a SUMO-Affimer and the other will not, functionally minimising the likelihood of steric clashes within this region. This construct, with the SUMO-Affimer within the MIR of the first HBcAg monomer within a fused dimer, is the first example of the VelcroVax system.

**Figure 1:**
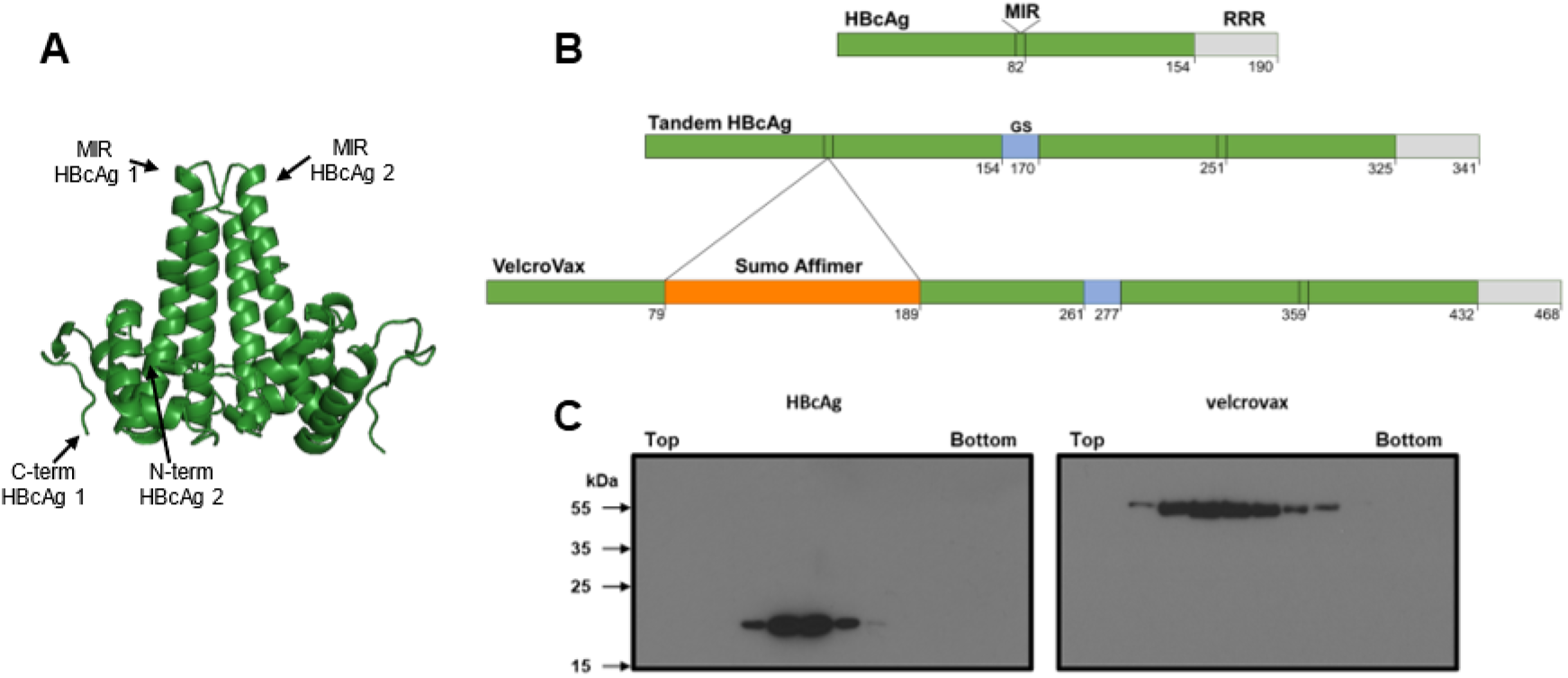
Generation of HBcAg and VelcroVax in Pichia pastoris. **(A)** X-ray crystal structure of a HBcAg dimer (PDB: 1QGT^48^). The locations of the C-terminal end of monomer 1 and N-terminal end of monomer 2, and the major immunodominant regions (MIR), are indicated. **(B)** Organisation of HBcAg, tandem HBcAg, and VelcroVax constructs with amino acid positions indicated. Representation depicts MIR, arginine-rich repeat (RRR), glycine-serine linking sequence (GS) and the insertion site of a SUMO Affimer within the MIR of the first of the fused HBcAg monomers. **(C)** Anti-HBcAg western blot of gradient purified HBcAg and VelcroVax particles produced in *Pichia pastoris*, probed with 10E11, representative figures, n = 3.

To determine whether the introduction of a SUMO-Affimer sequence within the MIR of a tandem HBcAg construct was compatible with particle formation, we utilised *P. pastoris* as an expression system. Samples of wt HBcAg or VelcroVax were produced in *P. pastoris* and separated along a 15-45% sucrose gradient (Fig 1C). Western blot analysis using anti-HBcAg antibody 10E11 indicated that both wt HBcAg and VelcroVax particles were present within the gradient. For both particle types, signal peaked around fraction 8, indicating that both wt HBcAg and VelcroVax effectively form VLPs in this system. Particle morphology was confirmed by negative stain EM (Fig S1).

### Structural characterisation of VelcroVax

To characterise VelcroVax structurally and assess the impact of SUMO-Affimer insertion, we generated high-resolution structures of VelcroVax VLPs. Notably, as a result of the tandem arrangement of VelcroVax, the *T* = 3* configuration does not conform to icosahedral symmetry. Each VelcroVax subunit comprises two connected HBcAg monomers, only one of which is modified with an Affimer, generating an imbalance between what would be true icosahedral asymmetric units (Fig S3A). As such, this configuration was termed *T* = 3* rather than *T* = 3, and five-fold (C5) symmetry was imposed during image processing rather than icosahedral symmetry (I1), which was imposed for the *T* = 4 VLP. Freshly purified VLPs were used for cryoEM data collection, and structures were determined for both *T* = 4 (at 2.9 Å resolution) and *T* = 3* (at 3.6 Å resolution) configurations (Fig 2, Fig S2).

**Figure 2:**
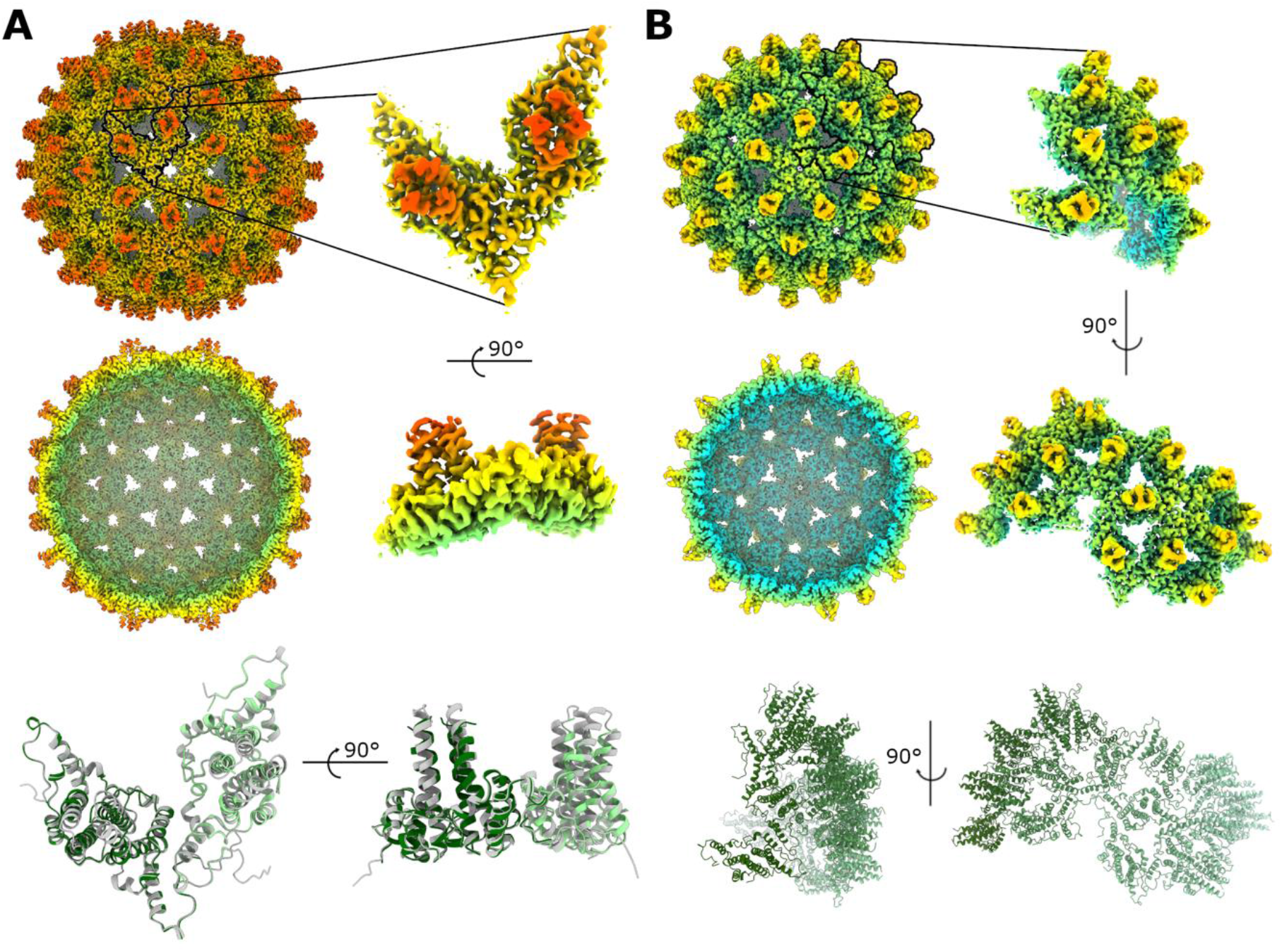
Structural characterisation of VelcroVax VLPs. Full and sectional isosurface representations of density maps for **(A)** *T* = 4 and **(B)** *T* = 3* VelcroVax VLPs, filtered by local resolution, shown at the same contour level and coloured according to the same radial colouring scheme. In each case an expanded view of an individual asymmetric unit (*T* = 4 – I1 symmetry; *T* = 3* – C5 symmetry) and corresponding atomic models are shown. For the *T* = 4 asymmetric unit, the VelcroVax atomic model (green) is overlaid with the cryoEM structure of wt HBcAg (grey, PDB: 7OD4^49^).

In general, VelcroVax showed a high level of structural similarity to unmodified HBcAg. For comparison, the atomic model for *T* = 4 VelcroVax was aligned with the best-matched subunit from a 2.8 Å resolution cryoEM structure of a *T* = 4 HBcAg VLP (PDB: 7OD4^49^). An RMSD value calculated between equivalent Cα atoms was only ~1.5 Å, and visual inspection revealed a high degree of overlap (Fig 2A). Most of the variation appeared to localise to the four-helix bundles, as might be expected given the proximity of this region to the inserted Affimer in VelcroVax.

Although the majority of the VLP was well resolved, density for the SUMO-Affimer was not evident in reconstructions of either *T = 4* or *T* = 3* VelcroVax VLPs. However, at low contour levels, weak, diffuse density was visible above four-helix bundles. For both *T* = 4 and *T* = 3* VLPs, maps low-pass filtered to 10 Å revealed additional density above the four-helix bundles consistent with the expected size of the Affimer (Fig 3), confirming that Affimers were likely present, but were not resolved to high resolution.

**Figure 3:**
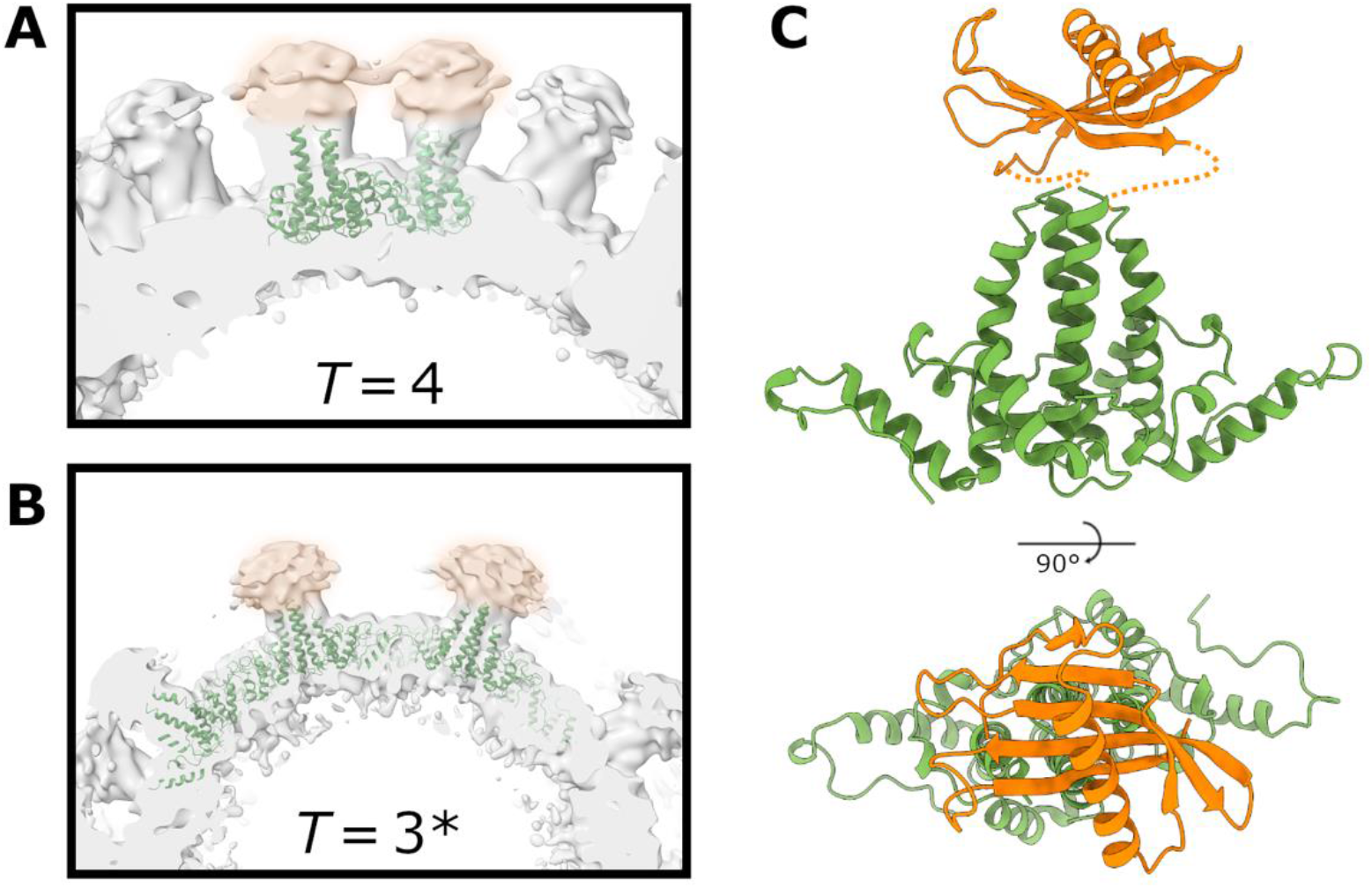
Affimer density in low-pass filtered VelcroVax VLP reconstructions. Sections of local resolution-filtered density maps for **(A)** *T* = 4 and **(B)** *T* = 3* VelcroVax VLPs following application of a 10-Å low-pass filter. Amorphous Affimer density (orange highlight) is visible above VelcroVax four-helix bundles. **(C)** Atomic model for a single VelcroVax monomer (green) with a SUMO-Affimer homology model (orange) manually positioned above the four-helix bundle, indicating the expected position of the Affimer based on the density shown in (A,B).

In an attempt to resolve Affimer density, data for the *T* = 4 configuration of VelcroVax was subjected to symmetry expansion and focussed 3D classification, using a mask to isolate the region above the four-helix bundle. However, while there was considerable variation between classes, none of the classes contained well-resolved Affimer density (Fig S4), confirming the high level of variability in Affimer positioning. Because of its unique symmetrical properties and therefore much more limited chance of success, focussed classification was not attempted for *T* = 3* data.

### Generation and capture of JUNV gp1

To determine whether VelcroVax retained a functional Affimer and thus was a suitable candidate for future immunisation work, we investigated the ability of VelcroVax particles to capture a SUMO-tagged antigen. Given its importance as a target for neutralising antibody responses^50–54^, we elected to use the gp1 subcomponent of the arenavirus gp1 spike from JUNV as a candidate immunogen. We firstly produced and purified C-terminally SUMO-tagged JUNV gp1 from HEK293T cells. The glycoprotein was purified with successive rounds of IMAC and SEC (Fig 4A), and SDS-PAGE followed by Coomassie blue staining verified the presence of glycoprotein within the peak fraction (Fig 4B). The binding of SUMO-tagged JUNV gp1 to VelcroVax was assessed by indirect ELISA. After coating EIA plates with PBS, wt HBcAg VLPs or VelcroVax overnight, wells were blocked, and glycoprotein was added. A JUNV gp1 specific antibody was used to detect the glycoprotein within each well. No JUNV gp1 was detected in the wells coated with PBS, or wt HBcAg. However, wells coated with VelcroVax bound JUNV gp1 in a concentration-dependent manner (Fig 4C).

**Figure 4:**
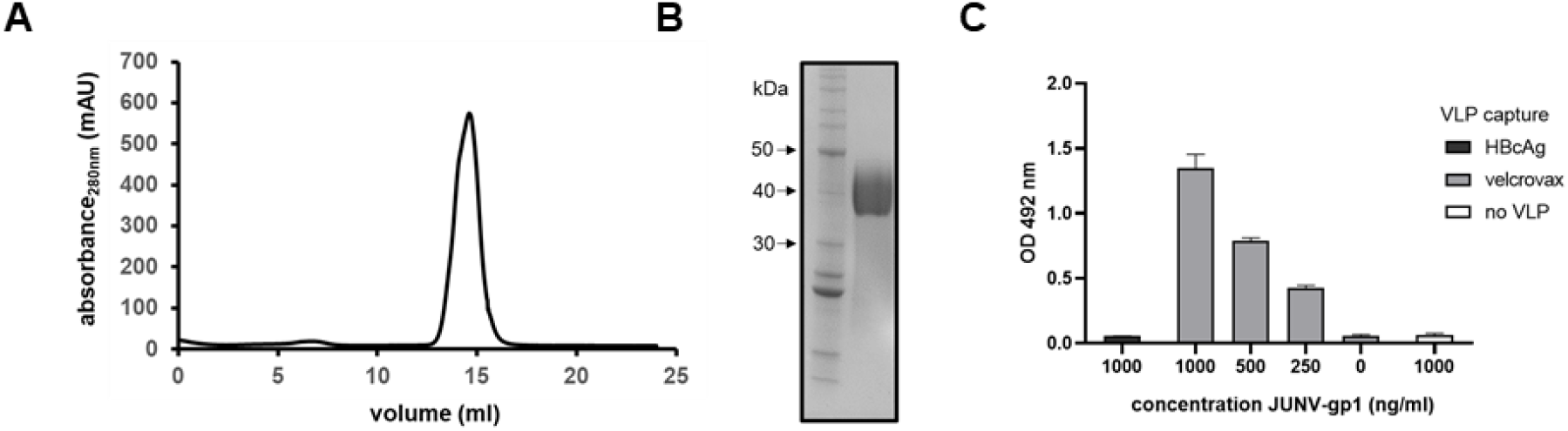
Generation of JUNV gp1 and interaction with VelcroVax. SUMO-tagged JUNV gp1 was produced in HEK293T cells and partially purified before processing through a final round of SEC. **(A)** Representative SEC elution profile for recombinantly derived JUNV gp1. **(B)** Reducing Coomassie-stained SDS-PAGE of SEC-purified JUNV gp1 with pertinent molecular mass standard sizes indicated in kDa. **(C)** ELISA was used to assess binding of HBcAg or VelcroVax to SUMO-tagged JUNV gp1. Particles coated on plates were subsequently incubated with JUNV gp1 and probed with anti-JUNV gp1 clone OD01-AA09, followed by incubation with anti-mouse HRP. Plates were incubated with OPD and the OD was read at 492 nm, graphed mean ± SEM, n = 3 in duplicate.

### Comparative immunisation

To compare the immunological consequences of immunisation with free glycoprotein with that presented on VLPs, immunisation trials were carried out in BALB/c mice. To this end, two groups of 7 mice were immunised three times at two-week intervals with JUNV gp1 mixed with wt HBcAg VLPs or bound to VelcroVax. Immunisations were administered subcutaneously in the presence of 2.5 nmol CpG ODN1668 and serum samples were collected between boosts, and two weeks after the final dose was administered (Fig S5).

Serum samples collected at completion of the immunisation series were assessed for the presence of IgG antibodies directed against HBcAg, VelcroVax, and JUNV gp1 (Fig 5A). Mice immunised with the wt HBcAg and JUNV gp1 generated antibodies reactive with HBcAg at titres greater than 1:4000. Although VelcroVax VLPs retain one unmodified HBcAg monomer per subunit, the antibodies generated against wt HBcAg recognised VelcroVax particles poorly. Similarly, the group immunised with wt HBcAg and JUNV gp1 did not generate high titre anti-gp1 antibodies. However, mice immunised with VelcroVax and JUNV gp1 generated antibodies which efficiently recognised JUNV gp1 and VelcroVax but not wt HBcAg (Fig 5a).

**Figure 5:**
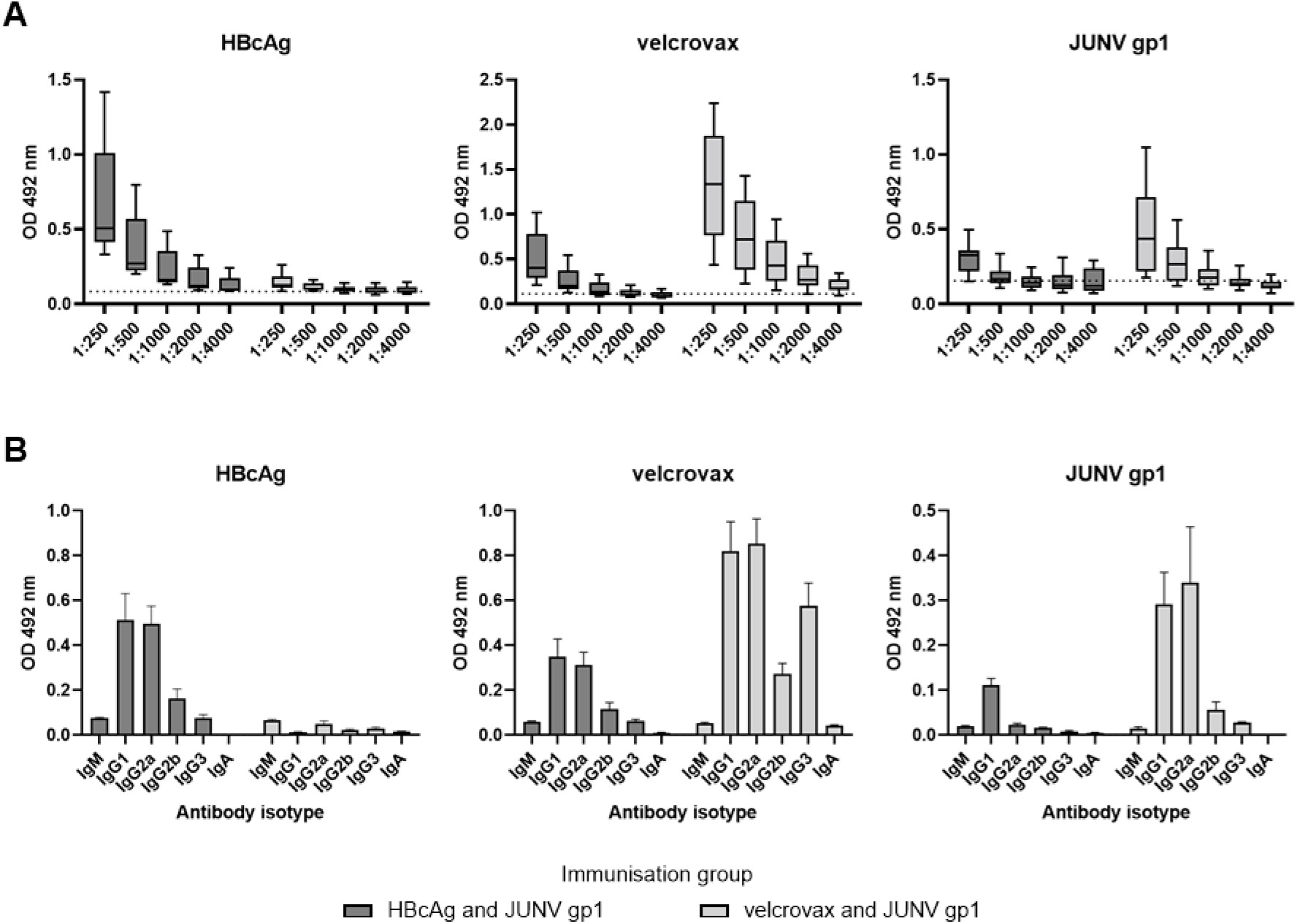
Reactive antibody titres and isotypes. Antisera generated by immunisation of mice with HBcAg and JUNV gp1 or VelcroVax and JUNV gp1 were assessed for **(A)** total reactive titres with HBcAg, VelcroVax, and JUNV gp1. Sera were assessed at dilutions between 1:250-4000, n = 7 in duplicate, graphed mean, 25^th^ and 75^th^ percentile with minimum and maximum ODs indicated. **(B)** Sera were subsequently assessed for isotype-specific reactivity with HBcAg, VelcroVax, and JUNV gp1. Sera were assessed at 1:125 dilution, n = 7 in duplicate, graphed mean and SEM.

To better understand the T helper (Th) bias of the immune responses generated we carried out antigen-specific isotyping of immune sera. Plates were coated with antigen and blocked as described above. Antisera were added to wells and incubated before the addition of isotype-specific detection antibodies. Both HBcAg and VelcroVax induced high levels of both IgG1 and IgG2a, suggesting a balance between Th1 and Th2 type responses (Fig 5B). Interestingly, despite the balanced response generated against HBcAg in the HBcAg immunisation group, the unbound JUNV gp1-specific antibodies induced were almost exclusively IgG1, indicating a strong Th2 bias directed against the glycoprotein. In contrast the anti-JUNV gp1 antibodies generated by VelcroVax-JUNV gp1 immunisation were balanced between IgG1 and IgG2a (Fig 5B), a potentially important characteristic for the development of effective viral vaccines.

We further assessed whether there was an isotype-specific bias in the responses against the peptide or glycan components of JUNV gp1. We incubated the glycoprotein with or without PNGaseF overnight before assessment by isotype-specific ELISA. While deglycosylation reduced the overall reactivity of antisera with JUNV gp1, there was no significant shift in the isotype preference (p = 0.3437) (Fig S6). No direct neutralisation of pseudovirus was detected using sera from either immunisation group at 1:100 dilution (Fig S7A). At a higher serum concentration (1:10) limited neutralisation was detected in some serum samples, and the VelcroVax group showed higher direct neutralisation at 1:10 dilution compared to the wt HBcAg immunisation group (P = 0.01), although mean neutralisation was just 24.89%. Additionally, neutralisation did not correlate with total reactive antibody titre (Fig S7B & S7C) or isotype (qualitative).

## Discussion

There is a global need for rapid development of vaccines that are adaptable to emerging pathogens and deliverable at low cost for use in LMICs. One approach to achieve this goal relies upon the development of a common carrier protein modified to present different haptens. Thus, a single carrier may be utilised as the foundation for vaccines against a range of pathogens, reducing vaccine development time and cost. To this end, we synthesised a carrier nanoparticle based upon the HBcAg protein, containing an adapter sequence to allow the post-purification coupling of haptens to VLPs. This nanoparticle forms the basis of a modifiable vaccine strategy. In addition to characterising the nanoparticle structurally, we selected JUNV gp1 as an exemplar hapten and determined the functional implications of hapten-VLP coupling on JUNV gp1 immunogenicity.

HBcAg VLPs are formed from monomers assembled into dimers, with 90 (*T* = 3) or 120 (*T* = 4) of these dimers assembling to form particles approximately 30 and 34 nm in diameter, respectively^55^. These particles are arranged with external facing N-termini, a long helical region followed by a flexible surface exposed loop (MIR), and another helical region leading to an internal facing C-terminal end (Fig 1A). The genetic fusion of two monomers results in a tandem HBcAg construct^56^, and the introduction of an anti-SUMO Affimer^27^ into the first MIR of this tandem construct forms the basis of our VLP capture system, VelcroVax (Fig 1B). Expression of HBcAg or VelcroVax in *P. pastoris* results in the efficient formation of VLPs, each having diameters consistent with the formation of both *T* = 3 and *T* = 4 symmetric particles (Fig 1C, Fig S1).

There are several published structures of wt and mutant HBcAg particles; however, no high-resolution structures exist of tandem HBcAg VLPs. Using VelcroVax particles produced in *P*. *pastoris,* we generated high-resolution structures of *T* = 3* and *T* = 4 symmetric particles, with the proportions of both particle configurations found to be approximately equal. The *T* = 3* reconstruction was less well resolved than the *T* = 4 reconstruction, likely because only five-fold symmetry was imposed during refinement to account for the unique symmetrical arrangement of *T* = 3* particles (Fig 2, Fig S2, Fig S3A). Both the *T =* 3* and *T* = 4 reconstructions had clearly resolved density for residues corresponding to the helices of both HBcAg molecules within the tandem VelcroVax sequence. Unsurprisingly, given the presence of flexible linking sequences, the SUMO-Affimer, the second MIR, and the internal Gly-Ser-linker lacked defined density. The fact that each fused dimer could occupy one of two orientations, leading to four unique arrangements per asymmetric unit for the *T* = 4 particle, also likely contributed to the poorly defined density of these regions (Fig S3B). Focussed classification yielded no improvement in Affimer density, and particles were distributed relatively evenly between focussed classes, suggesting a high level of variability and flexibility in this region, as expected (Fig S4). When a low-pass filter was applied to both *T* = 3* and *T* = 4 reconstructions, amorphous density was present above the four-helix bundles of the capsid, consistent with the presence of the Affimer (Fig 3). Given the difficulty in resolving flexible/mobile regions of the VLP at high resolution, we were unable to determine structurally whether Affimers displayed on the surface of particles retained a native fold.

Therefore, to determine whether Affimers expressed in the context of VLPs retained functionality we mixed SUMO-tagged JUNV gp1 with VelcroVax and assessed binding by ELISA (Fig 4). After confirming binding between VelcroVax VLPs and JUNV gp1, we carried out an immunisation trial using the complexed particles. The gp1 of JUNV forms a subcomponent of the trimeric gp spike displayed on the envelope surface and facilitates recognition of transferrin receptor 1 during host-cell entry^57,58^, suggesting it may be suitable for the generation of directly neutralising antibodies. Importantly, rabbits immunised with 3 doses of JUNV gp1 and adjuvant (80 μg/dose, GERBU Adjuvant P) generated 90% neutralisation at 1:20 serum dilution^59^. Another study suggested JUNV gp1 can elicit directly neutralising antibodies in mice, though this required three doses of 50 μg JUNV gp1 in the presence of complete Freund’s adjuvant^54^, which is approximately 50x the glycoprotein amount used here. Together these data suggest that antibodies directed against JUNV gp1 can be induced, though this does not appear to be particularly efficient and may require presentation as a part of the higher order gp. A commonality among JUNV neutralising antibodies is the presence of receptor-mimicking tyrosine residues (Ng et al, 2020). Interestingly, these tyrosine residues primarily arise in the CDRH3 region of neutralising antibodies, and these regions have the potential for greater diversity in species which utilise somatic gene conversion during antibody maturation^60^. Somatic gene conversion has been well documented in birds, sheep and rabbits; however, in humans and mice this mechanism has not been extensively reported^61^ and its relative role is unclear. Presentation of JUNV gp1 in the context of a nanoparticle vaccine may improve the maturation potential of antibodies directed against the glycoprotein subunit.

Nanoparticle vaccines are superior to isolated protein immunogens for several reasons. Their size (30-100 nm) facilitates improved recognition, uptake and enhanced antigen presentation^16,23,24^, and their repetitive structure enhances the crosslinking of receptors on B cells, which functionally improves signalling and is coupled with a shift in the cytokine milieu leading to a more balanced Th1/Th2 type response^26^. While the antibodies generated from immunisation only neutralised pseudovirus at low levels (Fig S7), the coupling of JUNV gp1 to VelcroVax both increased anti-JUNV gp1 antibody titres and generated a balanced Th1/Th2 response, as indicated by antibody isotypes (Fig 5). The stimulation of IgG2 antibodies has been associated with viral clearance in vaccination for influenza and thus the stimulation of Th1-type antibody may be desirable in vaccines seeking to limit disease severity^62^. As is the case with most peptide immunogens, this balance was not observed for the uncoupled JUNV gp1 immunisation group where anti-gp1 antibodies were predominantly IgG1 (Th2) (Fig 5b).

Similar to previous studies using JUNV gp1 as an immunogen, vaccination with JUNV gp1 coupled to VelcroVax failed to induce high-titre directly neutralising antibodies. Importantly, the broad response generated by the VelcroVax-JUNV gp1 complex indicates a more effective presentation of the target antigen when compared to unbound antigen. This broad response is generally desirable and, similar to responses directed against other viruses, may contribute to immunological protection in the absence of efficient direct neutralisation^62,63^. We therefore propose that the VelcroVax platform offers an adaptable system for future VLP vaccine applications.

## Acknowledgements

We would like to thank Shaun Baker for providing his expertise.

## Deposition of structural data

Atomic coordinates and density maps will be uploaded to the Protein Data Bank (PDB) and Electron Microscopy Data Bank (EMDB), respectively.

## Funding

We gratefully acknowledge support from The UK Medical Research Council MR/P022626/1 (NJK, JSS, NJS and DJR), MR/V031635/1 (TAB and KJD), and MR/S007555/1 (TAB). NJS, NJK and DJR are also supported by the NIH (R01 AI 169457-0). In addition, NJK and KG hold fellowships from Wellcome ISSF (204825/Z/16/Z) and JSS holds a Wellcome studentship (102174/B/13/Z). Electron microscopy was performed in the Astbury Biostructure Laboratory, University of Leeds, using equipment funded jointly by the University of Leeds and Wellcome (108466/Z/15/Z) and the Wellcome Centre for Human Genetics is supported by Wellcome Trust Core Award (203141/Z/16/Z).

## Author contributions

NJK, KG, NJS, DJR, MP, NJR, TAB conceived and planned experiments. Funding was sourced by NJS, NJK, DJR, KG and JSS. SS and AR generated the initial VelcroVax sequence, LWS and JA introduced this into *Pichia pastoris* and generated the material for structural studies. NJK generated the material for interaction and immunisation studies. NJK, KG and CRH performed ELISA to assess VelcroVax-JUNV gp1 interaction. TAB, GCP and AZ generated and purified JUNV gp1. DT selected the original SUMO-Affimer sequence. MH, MP and NJR carried out immunisation studies. KJD provided plasmids for JUNV pseudovirus production. JSS generated the structures of VelcroVax with the support of NAR. NJK determined antisera reactivity and isotypes. KG generated pseudovirus and carried out neutralisation assays. NJK prepared the initial manuscript with KG and JSS and all authors were involved in review of the data and editing of the manuscript.

## Supplementary material

**Figure S1:**
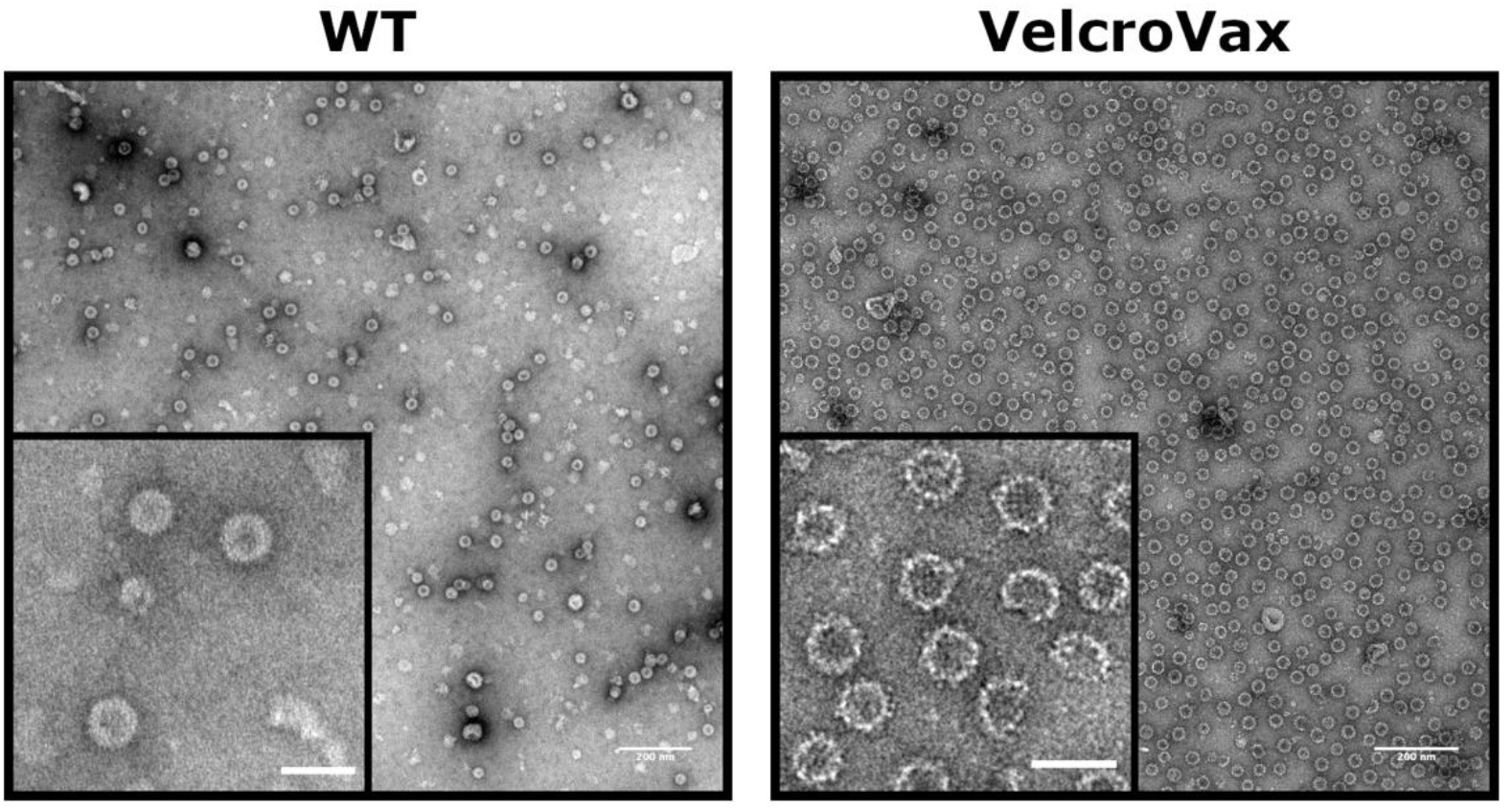
Characterisation of unmodified HBcAg and VelcroVax by negative stain EM. Representative micrographs of unmodified HBcAg (WT) and VelcroVax. For each, scale bars represent 200 nm (full micrograph) or 50 nm (expanded inset).

**Figure S2:**
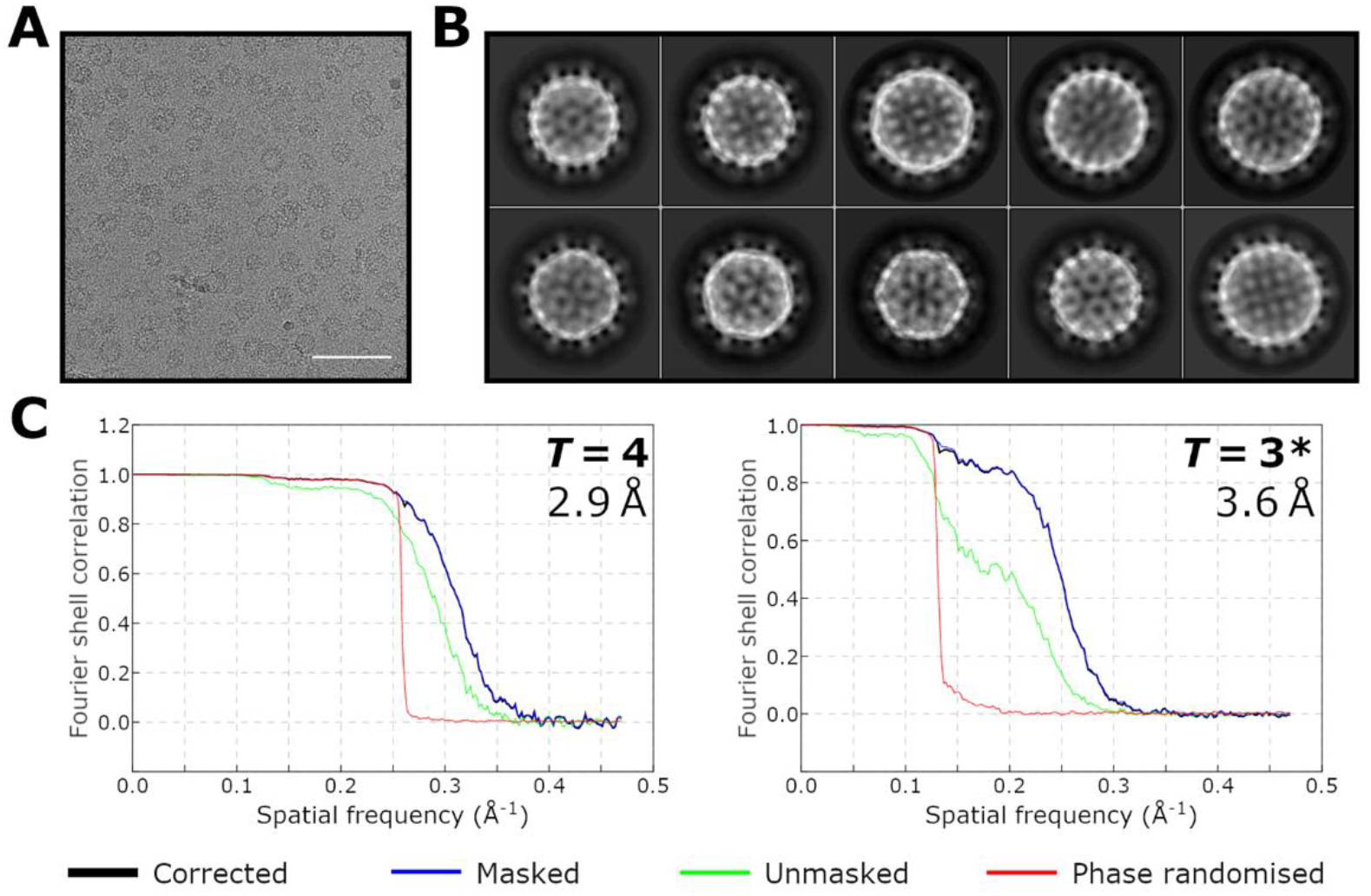
VelcroVax cryoEM data collection and image processing. **(A)** Representative micrograph from VelcroVax cryoEM dataset. Scale bar indicates 100 nm. **(B)** Representative class averages from 2D classification of VelcroVax particles, including both *T* = 4 and *T* = 3* VLPs. **(C)** Fourier shell correlation (FSC) plots for final reconstructions of *T* = 4 (left) and *T* = 3* (right) VLPs. Nominal resolutions are indicated, and were determined using the FSC = 0.143 criterion with high-resolution noise substitution to correct for any overfitting (black line, ‘corrected’).

**Figure S3:**
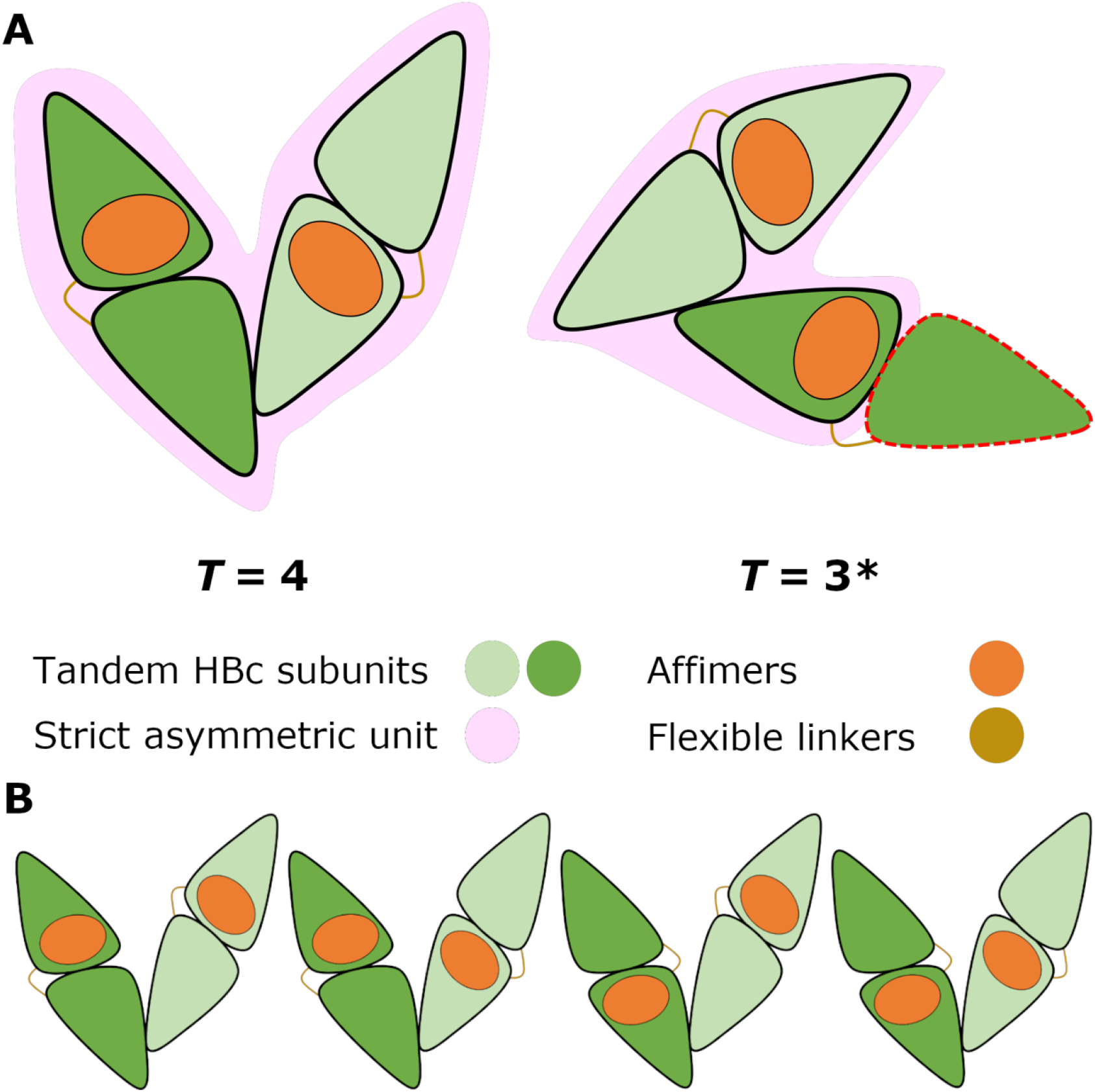
Inherent asymmetry within VelcroVax subunits. **(A)** Schematic illustrating how the tandem nature of VelcroVax does not conform to icosahedral symmetry in the *T* = 3* arrangement. Each VelcroVax monomer is formed from a tandem HBc subunit (green) linked by a flexible linker (beige) and a single Affimer (orange). This does not fit within the strict asymmetric unit (pink) of a true *T* = 3 VLP. **(B)** VelcroVax subunits can be incorporated into the asymmetric unit (here, *T* = 4) in either direction, leading to variation in the position of the Affimers. This results in blurring of Affimer density when particles are averaged to generate cryoEM reconstructions of VelcroVax VLPs.

**Figure S4:**
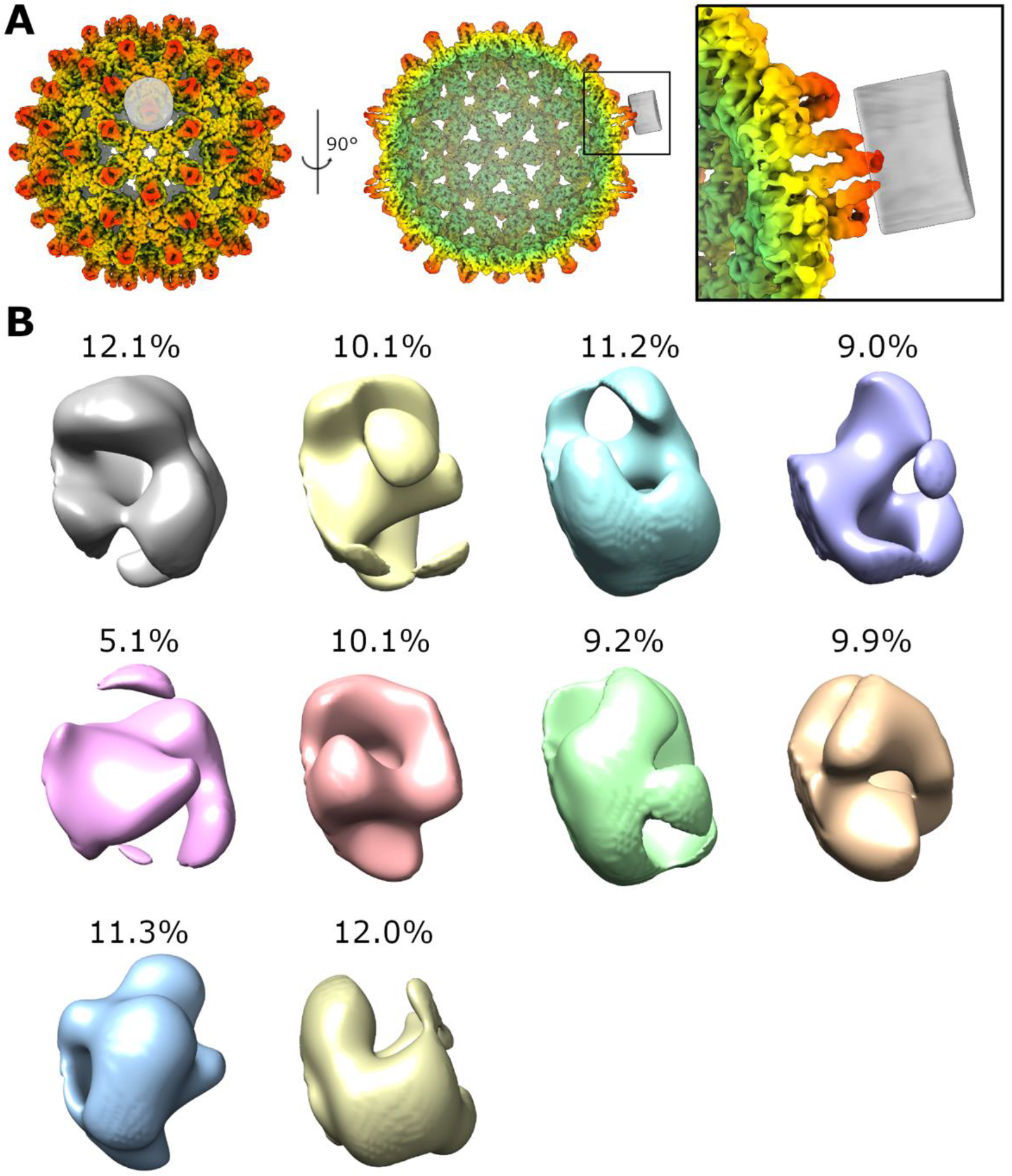
Focussed classification failed to resolve Affimer density. **(A)** Focussed classification was performed with a cylindrical mask (grey) positioned above a four-helix bundle from the reconstruction of VelcroVax in the *T* = 4 arrangement. **(B)** All classes from focussed classification, with the proportion of sub-particles assigned to each class indicated. Classes are shown oriented in the same way as the mask shown in the inset in (A).

**Figure S5:**
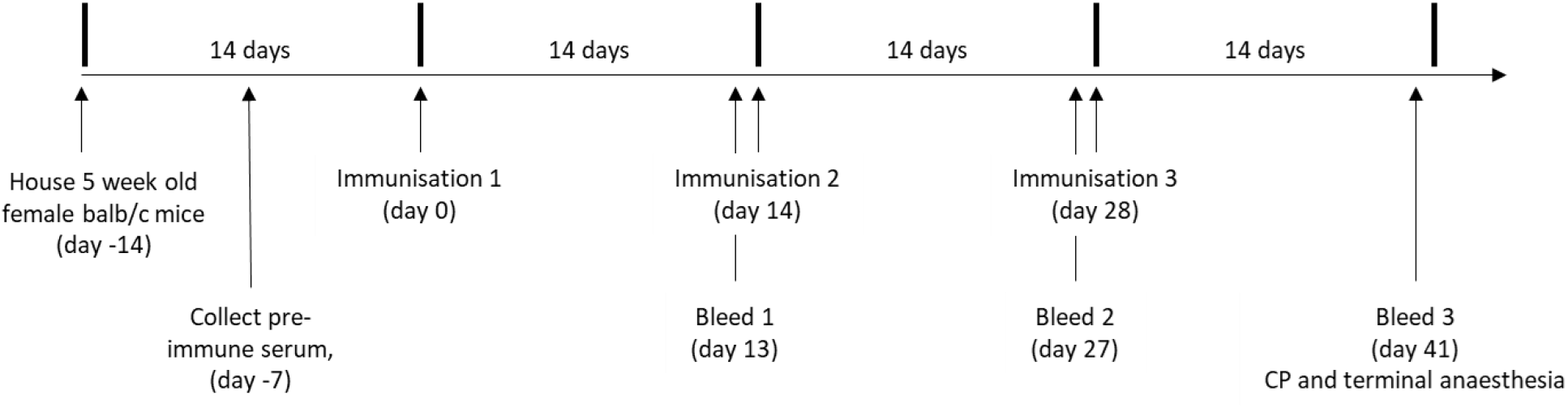
Immunisation schedule. Two groups of 7 female BALB/c mice were immunised three times at two-week intervals with a total of 2 μg protein, according to the above schedule. Vaccines were composed of 1 μg JUNV gp1, 1 μg VLP (HBcAg or VelcroVax), 2.5 nmol CpG ODN 1668 in a total volume of 200 μL. Intermittent blood samples were collected on day 13 and 27. At the conclusion of the experiment mice were humanely sacrificed, blood was collected via cardiac puncture while animals were under terminal anaesthesia.

**Figure S6:**
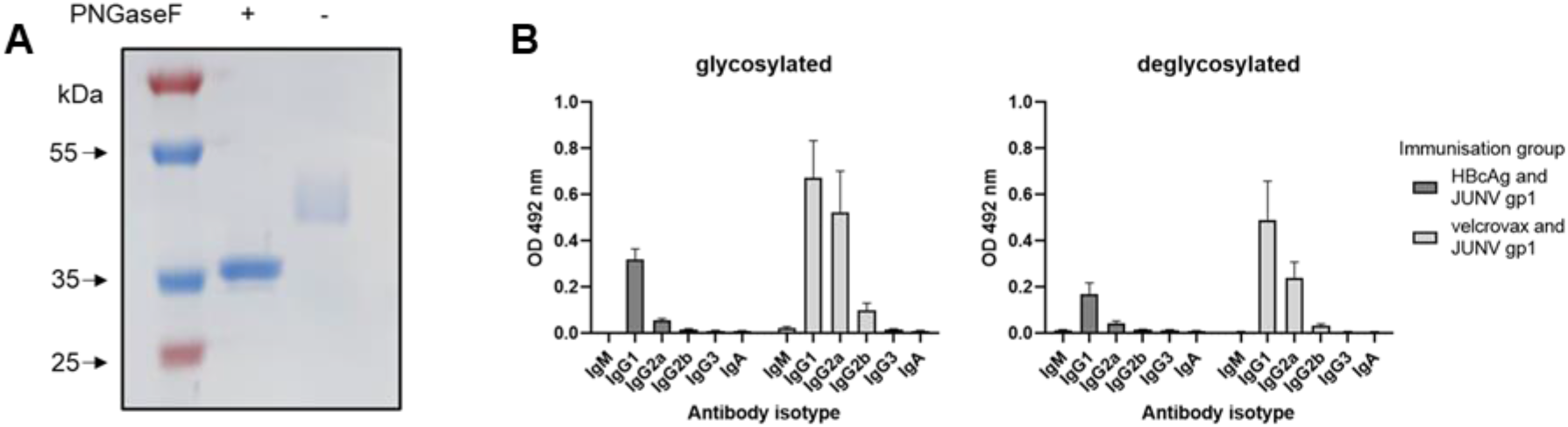
Glycan-specific antibody isotype. JUNV gp1 untreated or deglycosylated using PNGaseF. **(A)** Deglycosylation was confirmed using Coomassie stained SDS-PAGE. **(B)** Antisera generated from the immunisation of mice with HBcAg with JUNV gp1 or VelcroVax with JUNV gp1 were assessed for isotype-specific reactivity against glycosylated or deglycosylated JUNV gp1. Sera were assessed at 1:125 dilution, n = 7 in duplicate, graphed mean and SEM.

**Fig S7:**
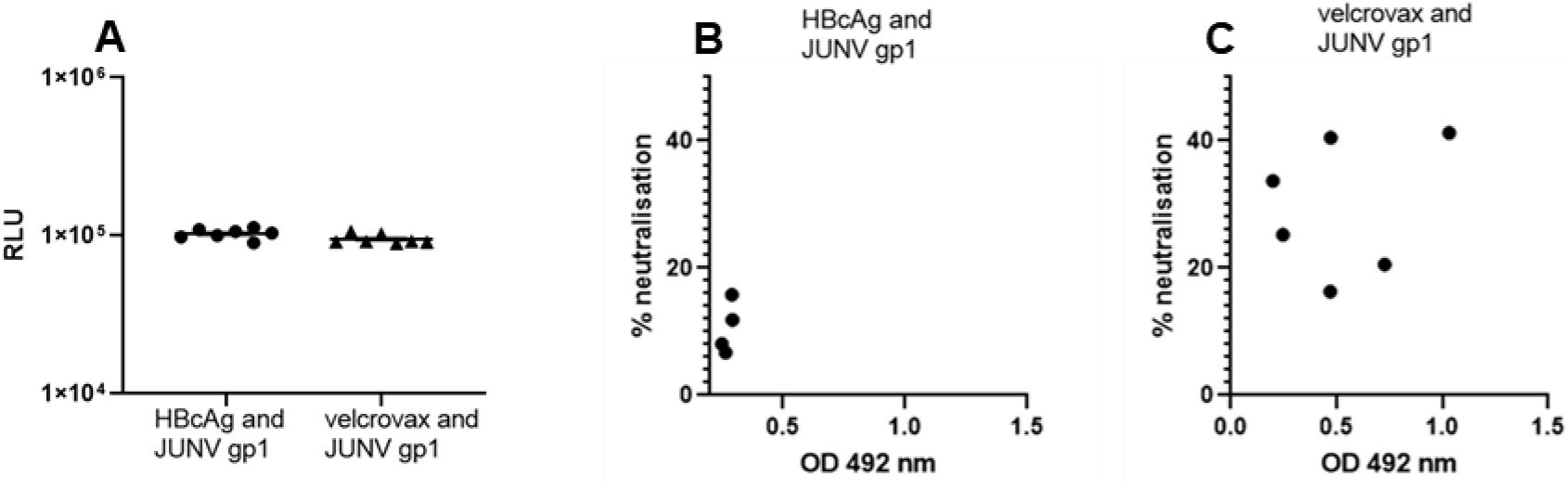
Pseudovirus neutralisation. JUNV pseudovirus was produced with a firefly luciferase reporter and used to transduce RD cells. **(A)** The ability of immune serum to directly neutralise 1×10^5^ RLU pseudovirus was assessed at 1:100 dilution. Data graphed showing average RLU of duplicate wells from individual animals, mean RLU/50 μL (n = 7) ±SEM. **(B and C)** Sera was tested for pseudovirus neutralisation at 1:10 dilution and graphed as % neutralisation relative to a non-serum containing control. Neutralisation from individual animals was graphed against total JUNV gp1 reactive titre at 1:250 dilution (complete reactive titres in Figure 5). Graphed mean values from duplicate pseudovirus neutralisation wells from individual animals, and mean OD 492 nm from n = 3 duplicate JUNV gp1 ELISA.

**Table S1:** CryoEM data collection parameters for VelcroVax.

|  | **VelcroVax** |
| --- | --- |
| **Microscope** | FEI Titan Krios |
| **Detector mode** | Linear |
| **Camera** | Falcon III |
| **Voltage (kV)** | 300 |
| **Pixel size (Å)** | 1.065 |
| **Nominal magnification** | 75,000× |
| **Exposure time (s)** | 1.3 |
| **Total dose (e<sup>-</sup>/Å<sup>2</sup>)** | 60 |
| **Number of fractions** | 40 |
| **Defocus range (μm)** | −0.8 to −3.0 |
| **Number of micrographs** | 3,643 |
| **Acquisition software** | Thermo Scientific EPU |

**Table S2:**
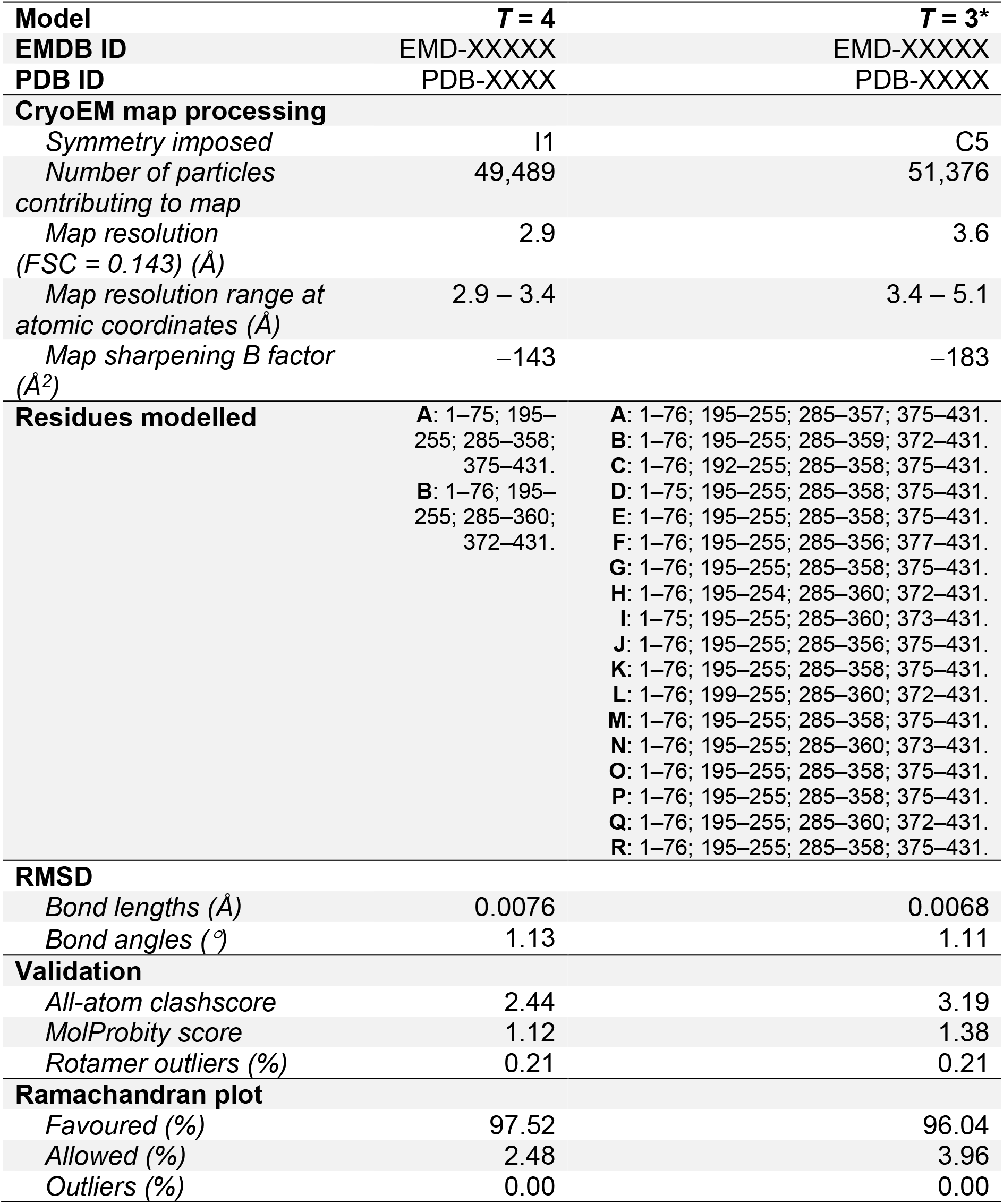
Quantitative parameters and validation statistics related to cryoEM image processing and model building.

| Model | *T* = 4 | *T* = 3* |
| --- | --- | --- |
| EMDB ID | EMD-XXXXX | EMD-XXXXX |
| PDB ID | PDB-XXXX | PDB-XXXX |
| **CryoEM map processing** |  |  |
| *Symmetry imposed* | I1 | C5 |
| *Number of particles contributing to map* | 49,489 | 51,376 |
| *Map resolution (FSC = 0.143) (Å)* | 2.9 | 3.6 |
| *Map resolution range at atomic coordinates (Å)* | 2.9 – 3.4 | 3.4 – 5.1 |
| *Map sharpening B factor (Å<sup>2</sup>)* | –143 | –183 |
| **Residues modelled** | A: 1–75; 195–255; 285–358; 375–431.<br>B: 1–76; 195–255; 285–360; 372–431. | A: 1–76; 195–255; 285–357; 375–431.<br>B: 1–76; 195–255; 285–359; 372–431.<br>C: 1–76; 192–255; 285–358; 375–431.<br>D: 1–75; 195–255; 285–358; 375–431.<br>E: 1–76; 195–255; 285–358; 375–431.<br>F: 1–76; 195–255; 285–356; 377–431.<br>G: 1–76; 195–255; 285–358; 375–431.<br>H: 1–76; 195–254; 285–360; 372–431.<br>I: 1–75; 195–255; 285–360; 373–431.<br>J: 1–76; 195–255; 285–356; 375–431.<br>K: 1–76; 195–255; 285–358; 375–431.<br>L: 1–76; 199–255; 285–360; 372–431.<br>M: 1–76; 195–255; 285–358; 375–431.<br>N: 1–76; 195–255; 285–360; 373–431.<br>O: 1–76; 195–255; 285–358; 375–431.<br>P: 1–76; 195–255; 285–358; 375–431.<br>Q: 1–76; 195–255; 285–360; 372–431.<br>R: 1–76; 195–255; 285–358; 375–431. |
| **RMSD** |  |  |
| *Bond lengths (Å)* | 0.0076 | 0.0068 |
| *Bond angles (°)* | 1.13 | 1.11 |
| **Validation** |  |  |
| *All-atom clashscore* | 2.44 | 3.19 |
| *MolProbity score* | 1.12 | 1.38 |
| *Rotamer outliers (%)* | 0.21 | 0.21 |
| **Ramachandran plot** |  |  |
| *Favoured (%)* | 97.52 | 96.04 |
| *Allowed (%)* | 2.48 | 3.96 |
| *Outliers (%)* | 0.00 | 0.00 |

